# One marine protected area is not enough: The trophic ecology of the broadnose sevengill shark (*Notorynchus cepedianus*) in the Southwest Atlantic

**DOI:** 10.1101/2023.01.25.524777

**Authors:** Manuela Funes, Agustín M. De Wysiecki, Nelson D. Bovcon, Andrés J. Jaureguizar, Alejo J. Irigoyen

**Affiliations:** Instituto de Investigaciones Marinas y Costeras (IIMyC), Consejo Nacional de Investigaciones Científicas y Técnicas (CONICET). Mar del Plata, Argentina; Centro para el Estudio de Sistemas Marinos (CESIMAR), Consejo Nacional de Investigaciones Científicas y Técnicas (CCT CENPAT–CONICET). Puerto Madryn, Argentina; Centro de Investigación y Transferencia Golfo San Jorge (CIT), Universidad de la Patagonia San Juan Bosco, Universidad Nacional de la Patagonia Austral, Consejo Nacional de Investigaciones Científicas y Técnicas (CONICET). Comodoro Rivadavia, Argentina; Instituto de Investigación en Hidrobiología, Facultad de Ciencias Naturales y Ciencias de la Salud, Universidad de la Patagonia San Juan Bosco. Trelew, Argentina; Comisión de Investigaciones Científicas de la Provincia de Buenos Aires (CIC). La Plata, Argentina; Instituto Argentino de Oceanografía (IADO). Bahía Blanca, Argentina; Universidad Provincial del Sudoeste (UPSO). Coronel Pringles, Argentina

**Keywords:** apex predator, diet, Patagonia, prey composition, stable isotope analysis, stomach content analysis

## Abstract

1. The broadnose sevengill shark (*Notorynchus cepedianus*) has been categorized as Vulnerable by the IUCN and shows a declining population trend in the Southwest Atlantic. Bycatch and poaching are the major threats in the region.
2. Although some ecological requirements have been described, there are still several information gaps regarding its ecology. Important aspects of its trophic ecology, like main prey items or key feeding grounds, remain uncertain and are essential to design effective conservation strategies.
3. We applied stable isotope and stomach content analyses to describe the trophic ecology of sevengill shark within a marine protected area of Peninsula Valdés in Patagonia, Argentina.
4. The stomach content analysis determined the southern elephant seal, *Mirounga leonina*, as the most frequently regurgitated prey item (70%*F*) during abundance peaks of both species in Península Valdes. The stable isotope analysis indicated that the overall contribution of the elephant seal to the diet of the sevengill shark was around 30% and that this percentage varied with the size of individuals.
5. Present results strengthen the current understanding of the trophic ecology of the sevengill shark. This study confirmed the use of the marine protected area as an essential foraging ground and identified its main prey items. Also, it reinforced the critical need to expand conservation tools beyond this particular coastal protection.

## 1. Introduction

Apex predators fulfill essential functions regarding ecosystem health (Estes et al., 2016;Fleming et al., 2016). They are considered sentinels of climate change (Hazen et al., 2019) and provide several ecosystem services, such as habitat engineering and nutrient cycling (Hammerschlag et al., 2019). Large sharks play key roles in the ecosystem structure and functioning of marine systems (Ferretti et al., 2010; Heithaus et al., 2008; Heupel et al., 2014;Roff et al., 2016). However, there is usually a lack of essential information on their ecology, which has been detrimental to the design of management strategies (Dulvy et al., 2014).

Shark vulnerability is conditioned by a combination of factors including habitat loss, pollution, climate change, and interaction with fisheries (Chin et al., 2010; Ferretti et al.,2010; Musick et al., 2000; Yan et al., 2021). Specifically, the broadnose sevengill shark *Notorynchus cepedianus* (Peron 1807) is currently categorized as Vulnerable by the International Union for the Conservation of Nature (Finucci et al., 2020). In the Southwest Atlantic Ocean (SWA), recent studies reported population declines of up to 60% (Barbini et al., 2015; Irigoyen and Trobbiani, 2016), and high bycatch rates within coastal and shelf areas with fishery exploitation (De Wysiecki et al., 2018; Góngora et al., 2009). In addition, there is a rising concern about poaching activities for fishery trophies in coastal areas (A. Irigoyen, pers. obs). One critical research need for shark conservation is to carefully address the resources which sustain the population and to identify its key feeding grounds (Simpfendorfer et al., 2011). This requires a broader understanding of its foraging ecology, main prey, trophic position, and intra-population patterns in the diet (Edwards, 2021; Heupel et al., 2014).

The sevengill shark has a wide latitudinal distribution in the SWA from southern Brazil (24°24’, De Wysiecki et al., 2020; Lopes et al., 2021) to the Magellan Strait in Chile (52°S, Guzmán M. and Campodónico G., 1976). During cold months, part of the population moves northwards across the SWA to Río de la Plata estuarine and Uruguayan coasts (De Wysiecki et al., 2020). During warmer months, part of the population moves southwards, probably related to foraging opportunities across coastal areas of Patagonia (De Wysiecki et al., 2020; Irigoyen et al., 2019, 2018). Particularly, in the protected area of Caleta Valdés, Península Valdes, the sevengill sharks are found year-round (Irigoyen et al., 2018). In this marine inlet, southern elephant seals *Mirounga leonina* also congregate in colonies to molt, reproduce, and rest (Campagna et al., 2000, 1998, 1998; Lewis et al., 2004). Between the austral spring and early summer (October to January), both elephant seals and sevengill sharks show abundance peaks in Caleta Valdés (Irigoyen et al., 2018; Lewis et al., 2004). During surveys in the area, spontaneous regurgitation of chunks of elephant seals consumed by sevengill sharks raised the hypothesis of Caleta Valdés as a key feeding ground (Irigoyen et al., 2018).

Studies that described the diet of sevengill sharks in the SWA reported elephant seals as a minor prey for the species (Crespi-Abril et al., 2003; Lucifora et al., 2005). These studies were performed using stomach content analysis, and because it reflects undigested items at the moment of capture, it only informs at short temporal and spatial scales (Baker et al.,2014). Considering the large-scale movements exerted by these migratory sharks, and therefore, the wide range of habitats used by them, there is a need for a diet description using a more integrative approach. In addition, the conservation status of this animal requires non-lethal techniques (Hammerschlag and Sulikowski, 2011). Stable isotope analysis (SIA) emerges as an opportunistic research tool because it integrates the diet information from broader temporal scales (Vander Zanden et al., 2015) and does not require animal sacrifice (Sanderson et al., 2009; Shiffman et al., 2012). This study aimed to provide a comprehensive description of the trophic ecology of the sevengill shark, identify the resources that support the population, and understand the importance of the aggregation site of Caleta Valdés as a feeding ground. We combined the information from regurgitations and SIA to identify its main prey items, determine its trophic level and investigate possible patterns in its diet.

## 2. Methods

### 2.1. Study site

The Caleta Valdés is an inlet with a complete marine regime located in northern Patagonia, Argentina (Fig. 1). The site is part of the Península Valdés marine protected area recognized in 1999 by UNESCO as a World Heritage Site. Although a priority has been placed to protect birds and marine mammals, it is also a spawning area for fish and crustacean species.

**Figure 1.**
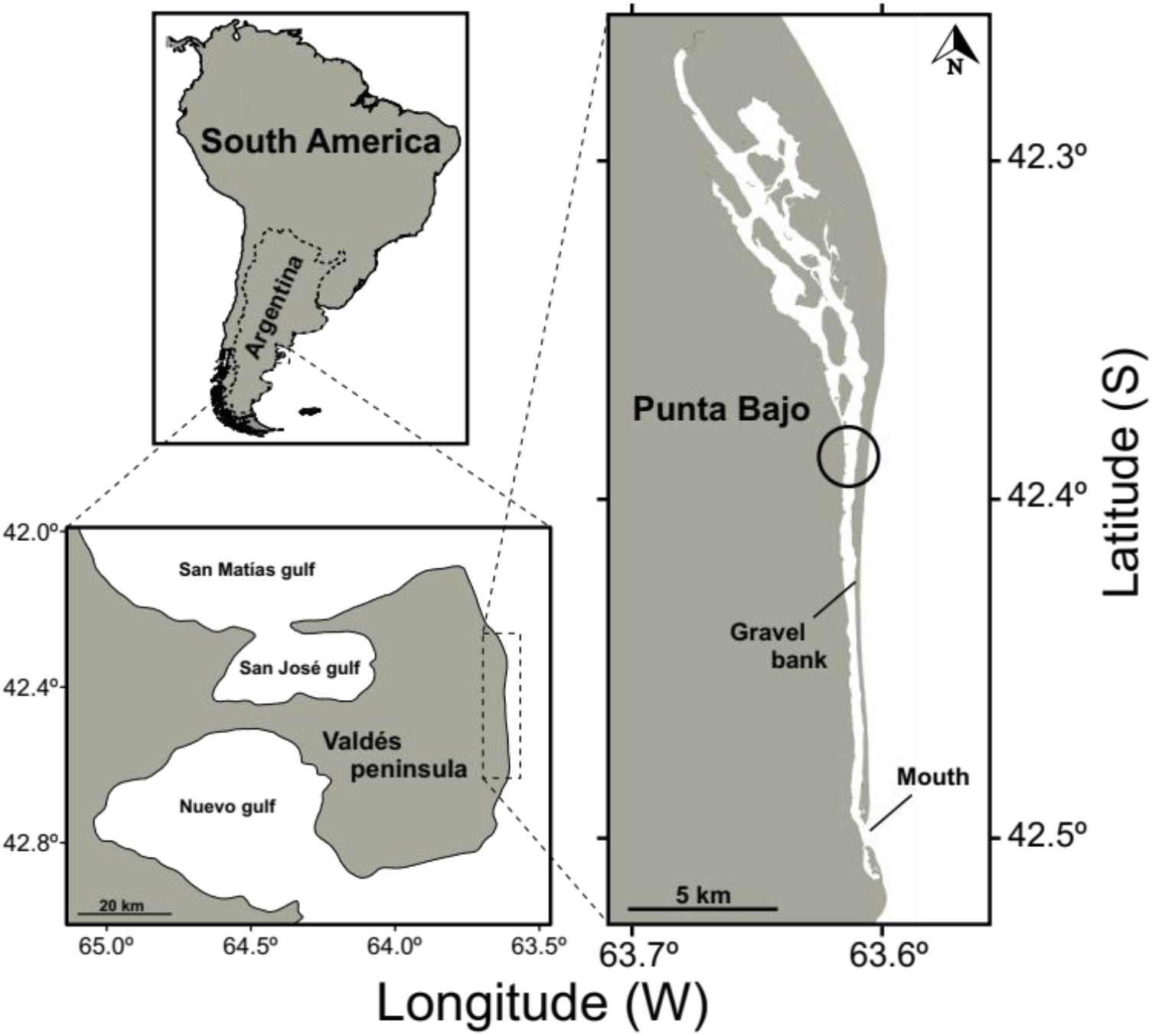
Study site location; Caleta Valdés within Península Valdés marine protected area, in northern Patagonia, Argentina.

### 2.2 Data collection and processing

As part of a bigger project studying the ecology of sevengill shark, sampling campaigns occurred every season between November 2015 and January 2017. Sharks were captured during standardized fishing sessions using longlines and rod and reel equipment.

Individuals were handled to the shoreline for sampling and safely released back into the water. Sharks were tagged on the dorsal fin with plastic spaghetti dart tags (floy-tags), sexed, and total length (*L*_T_) measured to the nearest cm. During handling procedures, several individuals spontaneously regurgitated stomach contents. All stomach contents were collected and identified to the lowest possible taxonomic level.

Blood samples (5 ml) were taken opportunistically by caudal venipuncture using syringes fitted with 14- or 18-gauge needles, depending on the size of the shark. Blood samples were transferred to plastic vials and stored at −20 °C. Samples were then dried at 60 °C until weight stabilized (~48 h) and grounded using a hand mortar. Carbon (δ^13^C) and nitrogen (δ^15^N) stable isotope compositions were determined by mass spectrometry at the Davis Stable Isotope Facility, University of California. Carbon and nitrogen stable isotope ratios were expressed in δ notation, in parts per thousand per mil (‰), with Vienna-Pee Dee Belemnite limestone and atmospheric N^2^ as standards for carbon and nitrogen, respectively (Peterson and Fry, 1987). The internal standards reported by the laboratory (mean and SD) were: G-13 bovine liver (δ^13^C = −21.7 ± 0.1 ‰, δ^15^N = 7.7 ± 0.1 ‰), G-18 Nylon 5 (δ^13^C = −27.7 ± 0.1 ‰, δ^15^N = −10.3 ± 0.1 ‰), G-20 glutamic acid (δ^13^C = −16.6 ± 0.1 ‰, δ^15^N = − 6.8 ± 0.1 ‰) and G-21 enriched alanine (δ^13^C = 43.0 ± 0.1 ‰, δ^15^N = 41.1 ± 0.1 ‰).

### 2.3 Data analyses

#### 2.3.1 Stomach contents

Prey composition in the stomach contents were determined as the frequency of occurrence (%F) of prey groups. This method was preferred over bulk composition because of its robustness and efficiency in the interpretation of diet composition (Baker et al., 2014).

#### 2.3.2 Mixing model analysis

The diet reconstruction of sevengill sharks was performed using mixing models in “MixSIAR” R package version 3.1.12 (Stock et al., 2018). This package uses Bayesian statistics to estimate the proportion of prey contribution to the consumer’s diet. The δ^13^C and δ^15^N values of the prey were taken from samples collected in the field and published stable isotope rates from the area (Table 1). To avoid exceeding the recommended maximum of sources (Phillips et al., 2014), prey items were grouped following ecological and isotopic value similarities. Elephant seals were represented in two different groups from distinct feeding grounds because these feeding grounds resulted in important differences in isotopic values (Eder et al., 2010). The trophic enrichment factors (TEFs) used were Δ^13^C = 1.7 ± 0.1 ‰ and Δ^15^N = 2.5 ± 0.2 ‰, which correspond to whole blood-specific values obtained from a controlled diet experiment with another elasmobranch species (Galván et al., 2016). A combination of sex and size were added as fixed factors to analyze possible patterns in the diet. The resulting models were compared through approximate leave-one-out cross-validation (LOO) (Vehtari et al., 2017). LOO estimates the probability that each model will make the best predictions on new data (Burnham and Anderson, 2002; McElreath, 2016) and is the most robust index when comparing Bayesian mixing models. Mixing models were run without concentration dependence (Phillips and Koch, 2002) and the error structure included the residual and the processing error terms (Parnell et al., 2013). Models were performed three times, running 500000 iterations and burning the first 100000; the convergence of the three Markov chain Monte Carlo was evaluated using Gelman and Rubin’s diagnostic (Stock et al., 2018).

**Table 1:** Stable isotope values of preys used to construct Mixing Models Analysis. Elasmobranchs and teleost values were estimated as the averages of values of the species contained in the group. Bibliographic sources and numbers of sampled individuals are also shown.

| Groups | Meand13C | SDd13C | Meand15N | SDd15N | N | Source |
| --- | --- | --- | --- | --- | --- | --- |
| **Elasmobranchs** | **-15.1** | **0.4** | **17.7** | **0.5** | **11** |  |
| *Squatina guggenheim* | -15.1 | 0.7 | 18.5 | 1.2 | 5 | Gaitán 2012 |
| *Atlantoraja castelnaui* | -15 | 0.3 | 17.6 | 0.6 | 3 | Gaitán 2012 |
| *Callorhynchus callorynchus* | -15.1 | 0.1 | 17.1 | 0.5 | 3 | Gaitán 2012 |
| **Teleosts** | **-18.1** | **0.8** | **17.3** | **0.4** | **23** |  |
| *Seriorella porosa* | -17.9 | 0.5 | 18.5 | 0.4 | 5 | Gaitán 2012 |
| *Merluccius hubbsi* | -17 | 0.5 | 17.1 | 0.4 | 10 | Drago et al. 2009 |
| *Stromateus brasiliensis* | -19.4 | 1.5 | 16.4 | 0.4 | 8 | Gaitán 2012 |
| **Sea Lions** | **-14.8** | **0.3** | **20.3** | **0.3** | **20** | Drago et al. 2009 |
| **Elephant Seals** |  |  |  |  |  |  |
| **Neritic ES** | **-15.3** | **0.5** | **17.1** | **1.4** | **4** | Eder et al., 2010 |
| **Oceanic ES** | **-18.8** | **0.4** | **13.1** | **0.45** | **9** | Eder et al., 2010 |

#### 2.3.3 Trophic position

The trophic position was estimated using the Bayesian modeling approach implemented in the R package “tRophicPosition” (Quezada-Romegialli et al., 2018). This model has the advantage of estimating trophic positions at the population level. The baseline was constructed with δ^13^C and δ^15^N values from six samples of zooplankton collected at the sample site with an inflated boat. The collection was performed using a net with a mesh of 180 μm and then split by size into two subsamples (<300 μm and >300 μm); it was assumed that the smaller fraction was mainly composed of small herbivores (Giménez, 2018; Pérez Seijas et al., 1987).

## 3. Results

### 3.1 Stomach Content Analysis

Over the course of the study, 496 sharks were caught. From the total, only 65 individuals spontaneously regurgitated stomach contents during handling (ten males and 55 females, 13.1% of total catch. Total size range 162–265 cm LT) (Fig. 2a). The majority of regurgitation events occurred in spring (*n* = 56), and nine in the rest of the seasons (two in autumn, three in winter, and four in summer, Fig 2b). The majority of individuals regurgitated southern elephant seals (*M. leonina*, *n* = 50, 77%*F*, usually soft pieces of muscle, fat, and skin, Fig. S1a). Other less frequent prey included silversides (*Odonthestes* sp., *n* = 6.1%*F*), patagonian blennie (*Eleginops maclovinus*, *n* = 3, 0.5%*F*), cockfish (*Callorhinchus callorynchus*, *n* = 4, 0.7%*F*), spiny dogfish (*Squalus acanthias*, *n* = 1, 0.1%*F*) and angel shark (*Squatina* sp., *n* = 1, 0.1%*F*) (Fig. S1 b-c; Table S1).

**Figure 2.**
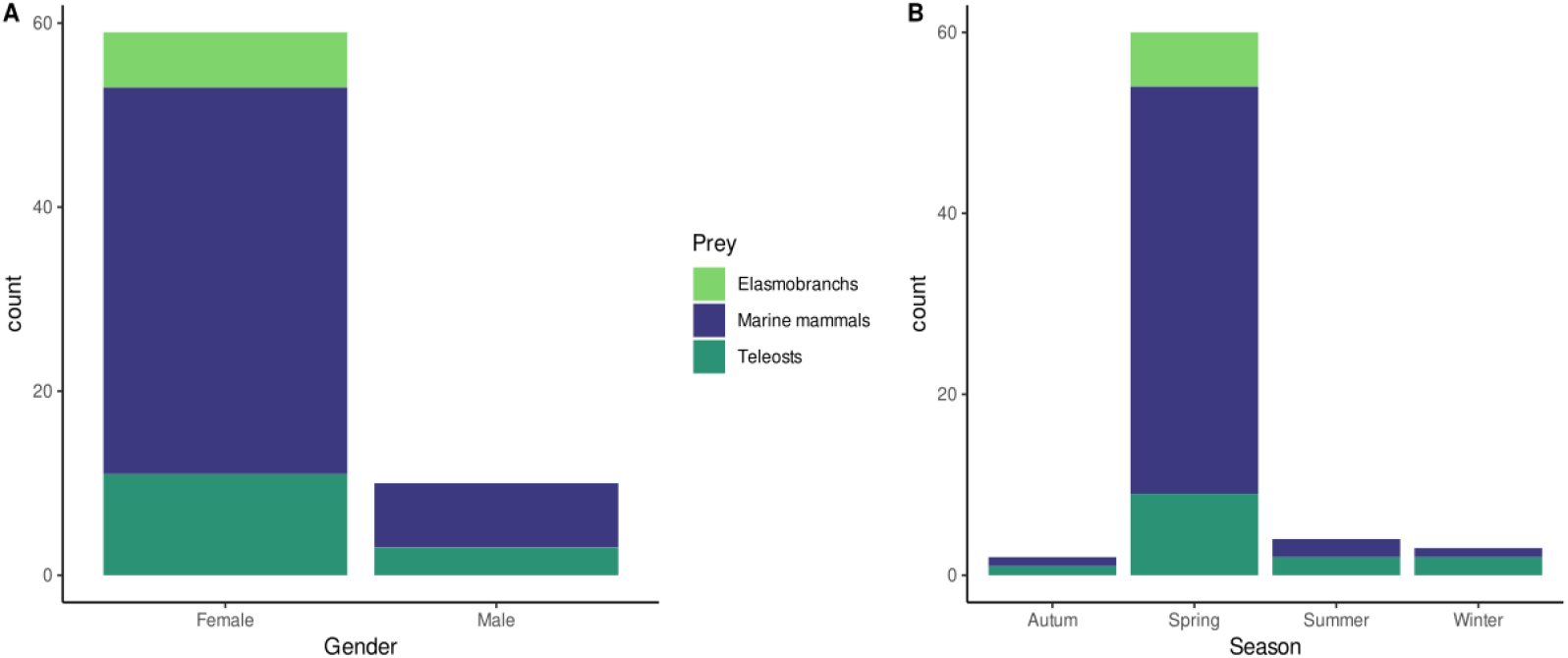
Stomach content analysis results of sevengill sharks (*Notorynchus cepedianus*) in Caleta Valdés. Results are expressed by the number of prey, color represents the group of prey. Panel **A** show results split by the sex of the shark and panel B by the season of occurrence.

### 3.2 Stable Isotope Analysis

A total of 58 blood samples (eight resamples, i.e. same individuals captured in different sampling events) were taken and analyzed from 50 sevengill sharks (30 females, 20 males; total size range 145–245 cm *L*_T_). Only eight individuals sampled for stomach content analysis were also sampled for stable isotope analysis. Eleven samples were collected in autumn, four in winter, 17 in spring, and 26 in summer. Isotopic values varied between 17.9‰ and 21.5‰ (mean = 20.1‰) for N and between −16‰ and −13.8‰ (mean = −14.7‰) for C. The calculated mean trophic position was 4.43 (CI_95%_ = 4.0–4.8).

The mixing polygon constructed with the selected prey (Table 1) satisfied the point in polygon assumption (Fig. 3a). The proposed mixing model showed differences in the isotopic compositions of the sources (Fig. 3a). The selected model included size as a continuous fixed factor (Table 2), which means that the relative contribution of each prey to the diet of sevengill shark varies with the size of the shark (Fig. 3b). The global mixing model showed that teleosts were the main contributors to sharks’ diet (mean = 37%; Table 3), whereas the combination of neritic and oceanic elephant seals was second in importance (19% and 10% respectively; Table 3). Elasmobranchs and sea lions were lesser contributors to the diet of the shark (17% and 13% respectively; Table 3). The consumption of neritic oceanic seals and elasmobranchs increased with size, whilst the consumption of the rest of the sources decreased with size (Fig. 3b).

**Figure 3.**
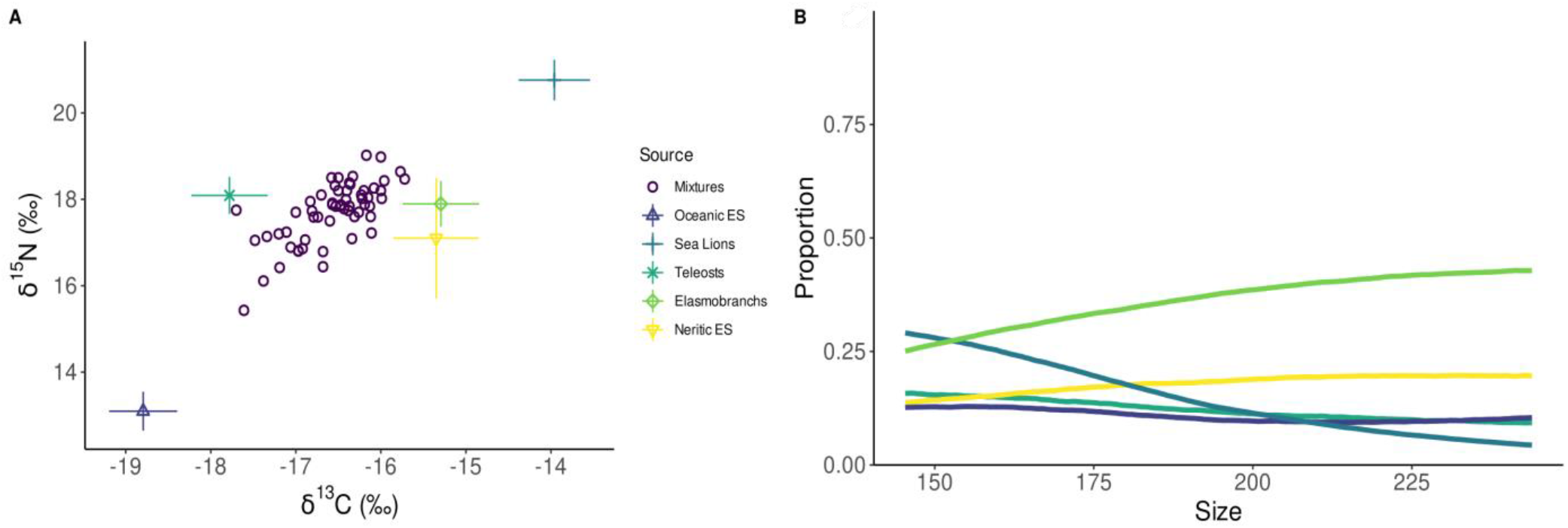
**A)** Mixing models δ^15^N and δ^13^C values of sevengill sharks (*Notorynchus cepedianus*) (violet circles) corrected by trophic enrichment factors from Galván et al. (2016). Mean and standard deviations (bars) of potential prey item values sampled in the SWA are indicated (Eder *et al.* 2019; Drago et al. 2009 & Gaitán 2012). **B)**Variation between the proportional contribution of each prey group to the diet of sharks and their size (total length).

**Table 2:** Comparison of mixing models fit using MixSIAR. Models are compared by approximate leave-one-out cross-validation (LOOic), standard deviation (se_LOOic), the difference in LOOic between each model and the model with the lowest LOOic (dLOOic), and the Akaike weight.

| Model | LOOic | se_LOOic | dLOOic | weight |
| --- | --- | --- | --- | --- |
| Size | 106.5 | 21.8 | 3.9 | 0.124 |
| Size+Sex | 102.6 | 21.6 | 0.0 | 0.871 |
| Null | 114.2 | 22.1 | 11.6 | 0.003 |
| Sex | 114.5 | 23.0 | 11.9 | 0.002 |

**Table 3:** Relative contribution of each source to the global diet of sevengill shark according to stable isotope values through MixSIAR analysis. Means, standard deviation, and 95% credibility intervals for each source are shown.

|  | Mean | SD | 95% |
| --- | --- | --- | --- |
| **Elasmobranchs** | 0.18 | 0.15 | 0.47 |
| **Neritic ES** | 0.20 | 0.14 | 0.43 |
| **Oceanic ES** | 0.11 | 0.07 | 0.25 |
| **Sea Lions** | 0.14 | 0.10 | 0.34 |
| **Teleosts** | 0.37 | 0.08 | 0.49 |

## 4. Discussion

This study was conducted on a species of conservation concern within a marine protected area of world heritage and under non-lethal techniques. This is also the first study applying SIA to the diet of the sevengill shark in the SWA. The combination of SIA and stomach content enabled an integrated description of the diet, which expands the current understanding of the feeding ecology of the sevengill shark. Also, we provide useful information on the importance of this marine protected area as a feeding ground for the sevengill shark.

Stomach content results showed that the elephant seal is a very important prey for the sevengill shark within the marine protected area. This preference for pinnipeds as prey was also reported in South Africa, where sevengill shark predates on South African fur seals (Braccini, 2008). However, does not match previuos descriptions in SWA, where the sevengill shark was characterized as generalist (Crespi-Abril et al., 2003; Lucifora et al.,2005). Although sampling efforts were balanced among seasons, stomach content results were unbalanced in terms of sex and season. Females were the most abundant gender, and most regurgitation events occurred in spring. Sevengill shark abundance in Caleta Valdes increase in warmer months, specifically in males (Irigoyen et al., 2018). In terms of prey, elephant seals are present in spring and summer but teleosts and elasmobranchs are present year-round. Therefore, the protective effect of the marine protected area may influence the diet preferences of sharks, i.e. when marine mammals are more abundant, they become the preferred prey.

In contrast with stomach content results, SIA revealed a more diverse diet and determined the teleosts as the main prey. The combined elephant seal groups contributed to almost 30% of the diet of sevengill sharks, highlighting their importance to the sevengill shark population. Considering that the SIA integrates diet information from several months (Logan and Lutcavage, 2010; Malpica-Cruz et al., 2012; Matich et al., 2017), the combined results agree with the importance of elephant seal consumption in Caleta Valdés.

The SIA results also showed that the diet varies with the size of the sharks. An increase in marine mammal consumption was previously described in relation to the increase in size (Lucifora et al., 2005; Ebert, 2002, Braccini, 2008). Present results showed an increase in neritic seals and elasmobranch consumption with increasing size, again, in agreement with what was reported before. However, results also showed a decrease in sea lion and oceanic seal consumption with increasing size. Differences with previous results could be explained by several factors, including the limitations of each sampling technique and the differences in sampling locations.

The estimated trophic position was 4.43, placing the species among the apex predators of the region (Ciancio et al., 2008). This species performs annual migrations (De Wysiecki et al., 2020) and feeds on mobile prey (Irigoyen et al., 2019, 2018; Lucifora et al., 2005). The sevengill shark is a powerful predator, it has a wide spectrum of possible prey and is capable of selecting prey according to their abundance or availability, which is common in sharks (Kim et al., 2012). Apex predator conservation has more importance on ecosystem conservation than previously perceived (Ferretti et al., 2010). Their vulnerability and possible indirect effects of trophic cascades have resulted in the widespread trophic downgrading of marine ecosystems (Baum and Worm, 2009; Estes et al., 2011).

Finally, this work highlights the role of coastal marine protected areas as a conservation tool for the sevengill shark population in the SWA. Evidence from the present study corroborates Caleta Valdés as an important foraging area for sevengill sharks. These areas offer transient protection during shark coastal aggregation events, in which they are particularly vulnerable to recreational fishers. Reasonable levels of protection within these marine protected areas are planned (e.g. permanent fishing ban in Caleta Valdés) however, illegal poaching is still a major issue in the area (A. Irigoyen, pers. observ., 2015 to 2021).

Investigating feeding habits and habitat use is crucial for understanding species-habitat relationships in the face of climate change (Birkmanis et al., 2020; Niella et al., 2022). Management strategies also require a holistic understanding of ecological demands to drive successful conservation actions that positively impact the stability of the ecosystems inhabited by sharks. In this sense, here we showed that sevengill sharks not only depend on protected resources but also on unprotected and harvested resources, located outside the marine protected areas. With the exception of the Buenos Aires Province, there are short or even null fisheries regulations for sharks in the remaining coastal jurisdictions of Argentina (Venerus and Cedrola, 2017). Considering its population decline, this work highlights the need to expand current conservation actions.

## Acknowledgments

We especially thank Silvestre Funes for his invaluable support and cooperation in the final phase of the work. To D. E. Galván for his time, ideas and help working with SIA data. To anglers G. Zamora, M. Cuestas, M. Lupiano, M. De Francesco, O. Trobbiani, and colleagues N. Sanchez-Carnero, F. Rios, M. Rossi, I. Rojo, J. M. Pereñíguez, R. Hernández, M. Pucheta and N. Demergassi for invaluable field support. To C. Awruch and A. Barnett for scientific advice. Funding was provided by the Shark Conservation Fund. Fieldwork included sampling within a World Natural Heritage Site and was authorized by Secretaría de Turismo y Áreas Protegidas del Chubut. MF, ADW and AJI were supported by the National Research Council (CESIMAR-CONICET, Argentina).

## Supplementary material

**Table S1.** Frequency of occurrence (%) of prey in stomach content analysis (*n* = 58) of sevengill sharks (*Notorynchus cepedianus*) during the peak of abundance (October-January) in Caleta Valdés, North Patagonia.

| Prey group | Regurgitations |
| --- | --- |
| **Teleosts** | **20.69** |
| *Bovichthys argentinus* |  |
| *Eleginops maclovinus* | 3.45 |
| *Engraulis anchoita* | 1.72 |
| *Odontesthes* sp. | 6.9 |
| *Pseudopercis semifasciata* |  |
| Unidentified rest | 8.62 |
| **Elasmobranchs** | **10.34** |
| *Callorhynchus callorhynchus* | 5.17 |
| *Squalus acanthias* | 1.72 |
| *Squatina* sp. | 1.72 |
| Rajidae | 1.72 |
| Unidentified rest | 3.45 |
| **Marine mammals** | **81.03** |
| Delphinidae | 5.17 |
| *Mirounga leonina* | 68.97 |
| Unidentified rest | 8.62 |

**Figure S1.**
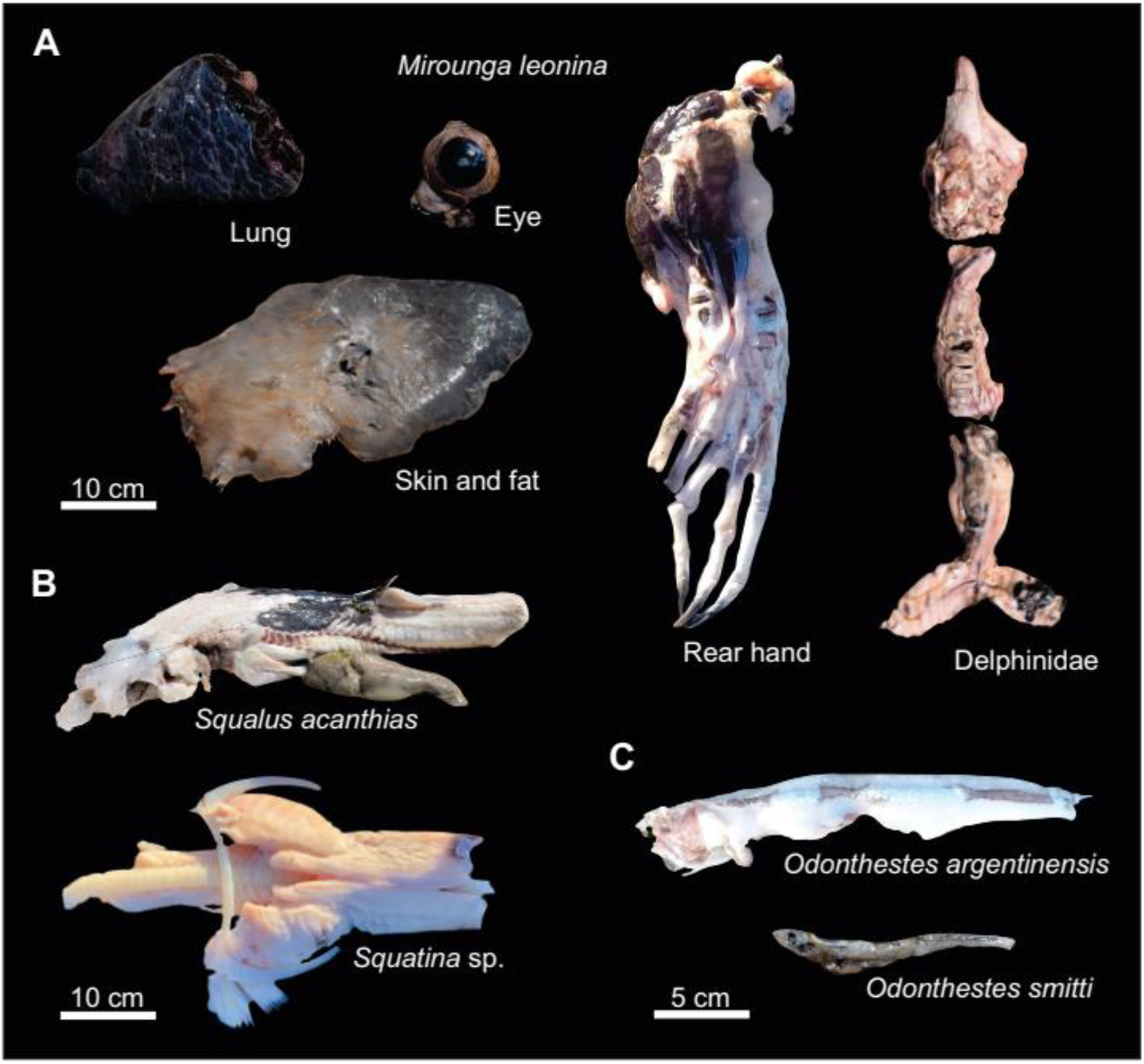
Photographic evidence of main regurgitated items from sevengill shark (*Notorynchus cepedianus*) individuals in Caleta Valdés. A-marine mammals, B-elasmobranchs, and C-teleosts.

